# First detection of peroxynitrite in live coral cells during thermal stress

**DOI:** 10.64898/2026.07.14.738561

**Authors:** Ioan D. Fuller, Kate P. Fetkenhour, G. Dinesh Kumar, Dylan W. Domaille, Liza M. Roger

**Affiliations:** School of Molecular Sciences, Arizona State University, Tempe, Arizona, USA; Department of Chemistry, Colorado School of Mines, Golden, CO, USA; School of Ocean Futures, Arizona State University, Tempe, Arizona, USA

**Keywords:** peroxynitrite, free radicals, reactive oxygen species, reactive nitrogen species, coral cell, oxidative stress, thermal stress, symbiosis

## Abstract

Reactive nitrogen species (RNS), particularly peroxynitrite generated from the reaction of superoxide and nitric oxide, are implicated in thermally-induced oxidative stress but remain difficult to resolve in live coral cells. We optimized fluorescent dye strategies to directly quantify superoxide, nitric oxide, and peroxynitrite production in thermally stressed *Pocillopora acuta* cell suspensions. Thermal stress was associated with an increase in intracellular peroxynitrite concentration, but not in its precursors, nitric oxide and superoxide, highlighting challenges with the application of fluorescent probes and their controls to live coral cells. Compounds developed for mammalian systems often translate poorly to non-model systems such as corals: strong endogenous fluorescence and multiple membrane barriers within the coral symbiocyte, for instance, limited the function of the nitric oxide probe, DAF-2DA. Despite these limitations, the detection of peroxynitrite in live, thermally stressed *P. acuta* cells represents a step forward in understanding the mechanism of coral bleaching. We also outline strategies for improving the performance of commercial dyes in non-model systems, including media optimization with EDTA treatment to preserve both cell viability and probe performance.

## Introduction

Coral reef ecosystems are among the most vulnerable to climate-driven change, and recent evidence suggests they may already lie beyond a critical threshold of resilience. A recent report identifies warm-water coral reef die-off as one of the five major Earth-system tipping points now at risk of being crossed under current warming trajectories (1). In this context, understanding the breakdown of the cnidarian-algal symbiosis that underpins reef productivity and structure takes on new urgency. Mechanisms of stress at the cellular and molecular scale now constitute a pivotal gateway to ecosystem-level collapse. Framing reef decline as both a symptom and a harbinger of global change, we turn to reactive oxygen and nitrogen species as a potential crucial mechanistic link between rising ocean heat and the rapid loss of reef function.

Oxidative stress is a side-effect of aerobic respiration, typically managed by endogenous antioxidant systems (2). Reactive oxygen species (ROS) and reactive nitrogen species (RNS) also play a crucial role in intracellular redox signaling (2). However, when produced in excess, these reactive species overwhelm the cell’s antioxidant capacity (3), which can lead to cell death (4,5). This redox imbalance is a central and unifying mechanism associated with coral bleaching (5–8), during which the mutualistic endosymbiosis between corals and dinoflagellates of the family Symbiodiniaceae collapses (7). Bleaching can be induced by a range of environmental factors, including temperature, salinity, irradiance, and pollution (8), but rising ocean temperature is the most pervasive and globally consequential driver today (7,9). As marine heatwaves intensify under climate change, understanding the cellular oxidative responses that precede bleaching is essential for predicting coral resilience and recovery.

The coral symbiocyte represents a complex redox microenvironment, comprising the host cytosol, the symbiosome lumen, and the algal endosymbiont cytoplasm, each characterized by distinct metabolic activities and oxygen gradients. Under optimal conditions, these compartments maintain a homeostatic balance of ROS and RNS production and scavenging. Thermal and light stress perturbs this equilibrium, disrupting photosynthetic and mitochondrial electron transport, and leading to ROS overproduction, including superoxide (O_2_•^-^) (10–12). Concurrently, host immune and signaling pathways elevate RNS, particularly nitric oxide (NO•) (13). A critical consequence of the simultaneous formation of superoxide and nitric oxide is the formation of peroxynitrite (ONOO^-^), a potent oxidant formed by the diffusion-limited reaction between the two (14). As such, peroxynitrite occupies a critical biochemical junction between nitrosative and oxidative stress (**Figure 1**), linking cellular signaling with oxidative damage. In animal and plant systems, peroxynitrite is recognized as a central mediator of redox-dependent pathology. It can nitrate tyrosine residues, oxidize lipids and nucleic acids, and modulate mitochondrial enzymes, thereby disrupting membrane integrity and energy metabolism (15). These molecular alterations can push the cell from reversible signaling responses toward irreversible injury. In the context of coral thermal stress, this reactivity suggests that peroxynitrite could represent a key inflection point, driving protein nitration, mitochondrial impairment, and ultimately symbiosis breakdown.

**Figure 1.**
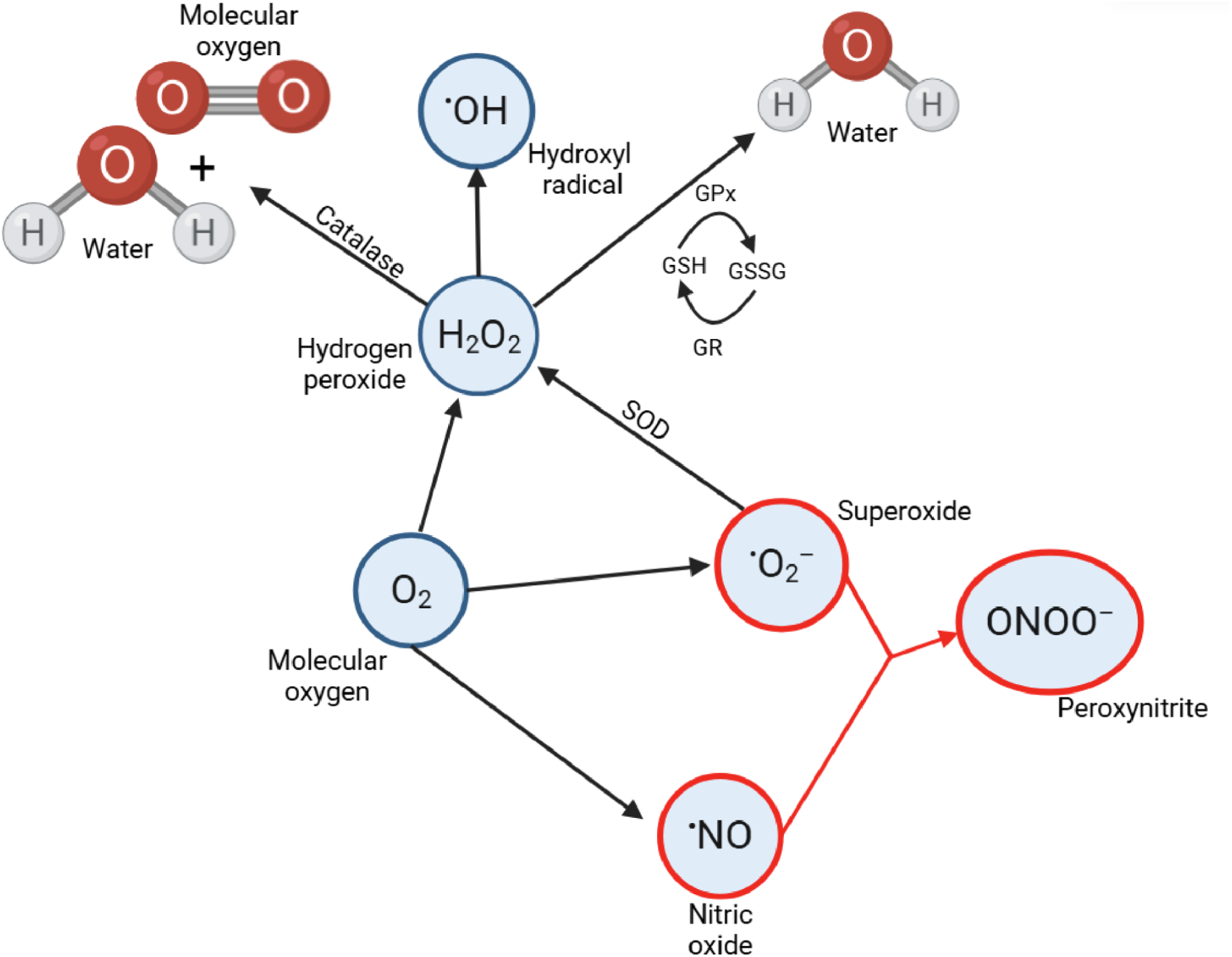
Diagram illustrating the interplay between oxidative and nitrosative pathways. Adapted from (16,17). Reactive species highlighted in red are the main components of nitrosative stress. Superoxide dismutase (SOD), glutathione peroxidase (GPx), glutathione (GSH), glutathione disulfide (GSSG), glutathione reductase (GR), and catalase are included to highlight how the antioxidant system works within the prooxidant system. Schematic created using BioRender

Despite extensive evidence implicating ROS and nitric oxide signaling in coral bleaching, the formation and role of peroxynitrite in coral cells remains largely unexplored: first implicated by Hawkins and Davy in *Exaiptasia* model anemone (18), it has since been characterized in coral tissue homogenate (19) but never in live coral cells. A larger body of work exists on the wider RNS pathway, documenting elevated nitric oxide production in cnidarians during thermal and light stress. This work has relied largely on model systems such as *Exaiptasia* (20–22), coral tissue homogenates (13,23) or free-living Symbiodiniaceae (24). Recent advances in coral cell culture (25) enable us to attempt direct detection of ROS and RNS within live coral cells, representing a significant step forward for the field.

To date, similar single-cell or intracellular analyses have relied solely on broad-spectrum fluorescent indicators such as CM-CH2DCFDA (26), which integrate the cumulative effects of multiple oxidants without chemical specificity. Nielsen et al. (26) reported no association between ROS production and physiological damage, and interpreted this as evidence against a role for ROS in bleaching. However, this conclusion rests on a broad-spectrum probe that sums the fluorescence of many oxidant species into a single signal. If damage is driven disproportionately by a minor species such as peroxynitrite, while the aggregate signal is dominated by an abundant, less-damaging species such as hydrogen peroxide, then total ROS output could remain flat or uncorrelated with damage even as the damaging species itself is fluctuating. A lack of association in the combined signal is therefore not sufficient to rule out a role for specific reactive species in driving cellular damage. We propose this as a testable hypothesis.

Applying techniques developed in mammalian model systems to coral cells presents several challenges. Probes used to detect nitric oxide, superoxide, and peroxynitrite were all developed and validated in mammalian systems (27–29), yet coral cells differ from mammalian cells in ways that can compromise probe performance: strong endogenous fluorescence (30) and the non-model buffer conditions required for coral cell culture (25) fall outside the pH and salinity ranges for which these probes were optimized. The structure of the coral symbiocyte compounds these challenges further. Unlike a typical mammalian cell, the symbiocyte comprises a plant cell, the algal endosymbiont, residing within an animal cell, the coral host, and separated from it by a symbiosome membrane (31). As a result, a fluorescent probe must cross three separate cell membranes and a plant cell wall to fully permeate the coral symbiocytes, rather than the single membrane typical of mammalian cell assays.

A number of ROS/RNS probes have naturally limited cell permeability and instead rely on acetate groups added to improve cell permeability: for instance, the nitric oxide probe, 4,5-diaminofluorescein diacetate (DAF-2DA) (27) and the general ROS probe, diacetyldichlorofluorescein (DCF-DA) (32). Once these probes are inside the cell, the acetate groups are cleaved by intracellular esterases, yielding the reactive, fluorescent product. However, this reactive product is commonly polar and therefore unable to cross further cell membranes (27). As a result, acetate-containing fluorescent probes are likely limited to detecting their target ROS/RNS within the cytosol of the coral host, rather than within the symbiosome or algal symbiont.

Mammalian cells thrive at pH 7.4 in low-salinity buffers such as 1X PBS (9‰) whereas seawater has a pH between 8.0 and 8.3 and a salinity of 35‰ (33). Most fluorescent probes are designed around a pH of 7.4, and their fluorescence intensity is commonly affected by environmental pH, including the nitric oxide probe DAF-2 (27) and the broad ROS probe DCFH-DA (34,35). Two probes used in this study are notable exceptions: the superoxide probe, MitoSOX Red, is reportedly stable up to pH 10 (36), and the far-red peroxynitrite probe exhibits constant response to peroxynitrite over pH 6–9 (29). The effect of salinity on fluorescent probe performance has been less widely studied; however, buffer selection has been shown to affect the performance of fluorescent probes, including a DAF-containing nitric oxide probe (37,38).

In this study, we translate dyes specific to the three key redox species implicated in coral bleaching: nitric oxide, superoxide and peroxynitrite, for use in a live coral system, along with a broad spectrum ROS dye for comparison. We use these dyes in live cell suspensions of *Pocillopora acuta*, a thermally sensitive species (39,40), subjected to conditions (35 °C) representative of a severe marine heatwave. This targeted approach is consistent with current best practice in the field (41). This study also serves as a preliminary investigation into the efficacy of commercially available fluorescent probes for imaging ROS and RNS in a live non-mammalian system, i.e., the coral symbiocyte (coral host with algal symbiont).

## Materials and Methods

### Materials

Filtered sterile artificial seawater (FASW) for use in cell culture was prepared using the salt composition described by (33). Briefly, solutions of hydrous salts, anhydrous salts, and sodium carbonate were prepared in Milli-Q water (18.2 MΩ), sterilized separately using an autoclave and allowed to rest (48 h) before combining the solutions. Salinity was adjusted to 35‰ using sterile Milli-Q water and the resulting ASW was filtered using a 0.22 μm filter (Millipore Express® PLUS, Merck, IRL).

Cell culture media (CCM) contained FASW (74% v/v), Dulbecco’s Modified Eagle Medium (15% v/v), Fetal Bovine Serum (10% v/v), Antibiotic-Antimycotic (0.5% v/v), gentamicin (0.5% v/v). CCM was prepared weekly.

EDTA-containing artificial seawater (ASW with EDTA) was prepared by adding EDTA to FASW to a final concentration (f.c.) of 20 μM.

Calcium-magnesium-free artificial seawater was prepared as by (Roger & Chowdhury, 2026). Briefly, salts were added to deionized water, autoclaved and filtered (0.22 μm; Millipore Express® PLUS, Merck, IRL).

Dyes, positive and negative controls for assays are listed here: 4,5-diaminofluorescein diacetate (DAF-2DA); luteolin (LUT); sodium nitroprusside (SNP); BioTracker Far-Red Peroxynitrite Live Cell Dye (FRD); Ebselen (EBS); 3-Morpholinosydnonimine hydrochloride (SIN-1); MitoSOX Mitochondrial Superoxide Indicator Red (MitoSOX); Superoxide dismutase (SOD); MitoTEMPO (MTPO); 6-carboxy-2’,7’-dichlorodihydrofluorescein diacetate, di(acetoxymethyl ester) (carboxy DCFDA-AM); hydrogen peroxide; PNP-1. Details about each material are compiled in **Table S1**.

### Cell dissociation and culture

Cell dissociation was performed as previously described (42). Briefly, a *P. acuta* colony (husbandry details are provided in **S1**) was placed in a crystallization dish containing ASW with Coral Rx disinfectant (25 µL/mL ASW), nubbins (∼1 cm each) were cut from the colony with sterile clippers and allowed to rest with constant bubbling for 10 min. The nubbins were transferred to a biosafety cabinet and each nubbin was rinsed with FASW (∼1 min) to remove any skeletal debris and mucus before incubation in calcium-magnesium-free ASW (3 mL/nubbin) for 1 h under ambient light. Post incubation, each nubbin was rinsed with calcium-magnesium-free ASW, sloughing cells off the skeleton to form a cell suspension. The resulting cell suspensions were combined and centrifuged (805 × *g*, 3 min, 25 °C). The supernatant was discarded and the cell pellet resuspended in CCM (0.5 mL/nubbin). Cell viability was performed as previously described (30), and is detailed in **S2**.

### Assay preparation: imaging

Samples for assays were prepared using four different fluorescent probes: DAF-2DA (nitric oxide-specific, intracellular) with SNP positive control (21) and LUT negative control (43,44); MitoSOX Red (superoxide-specific, intracellular) with SIN-1 positive control, SOD negative control (22), and MTPO negative control (45,46); FRD (peroxynitrite-specific, intracellular) with SIN-1 positive control and ebselen negative control (29); carboxy-DCFDA-AM (general ROS). All cell suspensions were diluted to 1 × 10^6^ live cells/mL in ASW with EDTA. Each experiment included four treatments: untreated *P. acuta* cell suspension; *P. acuta* cell suspension treated with probe (unknown); *P. acuta* cell suspension with probe and positive control; *P. acuta* cell suspension with probe and negative control. Detail on each treatment for each dye is provided in **S3**–**S6.** Each treatment was performed under both control (25 °C, 1 h) and thermal stress (35 °C, 1 h) conditions. All samples were imaged in a chambered slide on an ECHO Revolve microscope. Micrographs were captured using ECHO software at 100× magnification with oil immersion under optimized intensity settings for the following fluorescence cubes: DAPI (λex: 380/30 nm, λem: 450/50 nm) 100% intensity, 20.0 ms exposure, high gain; TRITC (λex: 530/40 nm, λem: 605/70 nm) 49% intensity, 10.0 ms exposure, high gain; FITC (λex: 470/40 nm, λem: 525/50 nm) 38% intensity, 15.0 ms exposure, high gain; CY5 (λex: 630/40 nm, λem: 700/75 nm) 49% intensity, 10.0 ms exposure, high gain); along with transmitted light and bright field.

### Assay preparation: time-series

Thermal stress conditions were selected to reflect a graded severity of heat stress: 33 °C was used as an estimated sublethal thermal stress threshold for *P. acuta*, while 35 °C was used to represent more severe marine heatwave conditions and to test whether reactive species signals intensified with increasing thermal stress severity. Three fluorescent probes were trialled within this system: DAF-2DA (for nitric oxide), MitoSOX Red (for superoxide) and carboxy-DCFDA-AM (for broad spectrum ROS). Hydrogen peroxide was used as a positive control, and dye and cell suspension blanks were included with each experiment. Full details can be found in **S7**–**S9**.

### Assay optimization

The influence of buffer selection on fluorescent dye performance was assessed using a peroxynitrite-specific dye, PNP-1 (47) in four buffer conditions: 1X PBS (diluted from 10X PBS pH 7.4), 1X PBS (pH 8.3), CCM (pH ∼8.3), ASW+EDTA (pH ∼8.3). Where necessary, buffer pH was adjusted by adding dropwise amounts of sodium hydroxide (25 mM) and change monitored using pH indicator strips (pH 6.5–10.0).

PNP-1 (f.c. 4.9 µM) was added to each buffer in a black 96-well plate. Peroxynitrite (f.c. 719 µM and 7 µM) was added to each buffer to test for sensitivity. Hydrogen peroxide was added to each buffer (f.c. 87 mM) to test for selectivity. Control wells included only buffer and PNP-1. The plate was incubated (25 °C, 30 min, dark) in a plate reader before measuring fluorescence spectra (λex/em = 450/500–650 nm, Δλem = 1 nm). The average fluorescence of each buffer plus PNP-1 control was subtracted from the peroxynitrite and hydrogen peroxide-containing wells to correct for background fluorescence at each wavelength.

### Data

Intensity values were obtained from micrographs using ECHO software: regions of interest (ROIs) were drawn in the cells and the values extracted as .csv files. For all microscopy trials except carboxy-DCFDA-AM and MitoSOX positive control experiments, 15 micrographs were analyzed in total, five images from each experimental replicate. For carboxy-DCFDA-AM, 10 micrographs were analyzed from one experimental replicate. For the MitoSOX positive control experiment, 10 micrographs were analyzed from one experimental replicate.

Values were organized by temperature and dye treatment, and plotted in R. Analysis of data was performed using R 4.5.2 where statistical analyses were performed (ANOVA, Tukey, paired t-test, Wilcox test).

Fluorescence intensities were exported from the microplate reader using Agilent BioTek Gen5 software in .xlsx format, processed and analyzed using Microsoft Excel and R 4.5.2. Each experiment included 12 wells with *P. acuta* cells stained with fluorescent probe, 3 wells with a hydrogen peroxide-spiked control, 6 wells with a dye-only control and 8 wells with unstained *P. acuta* cells.

## Results

Cell viability as assessed post-dissociation, prior to experiments; the initial viability of dissociated *P. acuta* cells used for these experiments was 92 ± 1.3%. The effect of buffer on fluorescent probe performance was tested with the peroxynitrite-specific probe, PNP-1 (47); we found that ASW with EDTA (20 µM) offered the best balance of probe performance while maintaining salinity and pH appropriate for coral cells (**Figure S1**) cell viability. All imaging trials were performed using ASW with EDTA, while the time-series trials were performed with CCM, to maintain cell viability over the extended incubation period.

### Peroxynitrite

The average relative fluorescence intensity in cells loaded with the peroxynitrite dye FRD was 324.89 ± 13.15 RFU at 25 °C compared to 471.49 ± 8.46 RFU at 35 °C (Figure 2A), a significant increase (p < 0.05, Tukey’s test) attributable to applied thermal stress and likely corresponding to an increase in peroxynitrite production. Intracellular FRD fluorescence is shown in **Figure 3**. The average relative fluorescence intensity of SIN-1 (positive control) was 416.59 ± 8.07 and 384.02 ± 7.15 RFU for 25 °C and 35 °C respectively; no significant difference was found between this positive control and our unknown (i.e. cells with dye), suggesting that SIN-1 was not an effective source of nitric oxide and superoxide (the two precursors to peroxynitrite) in this system.

**Figure 2.**
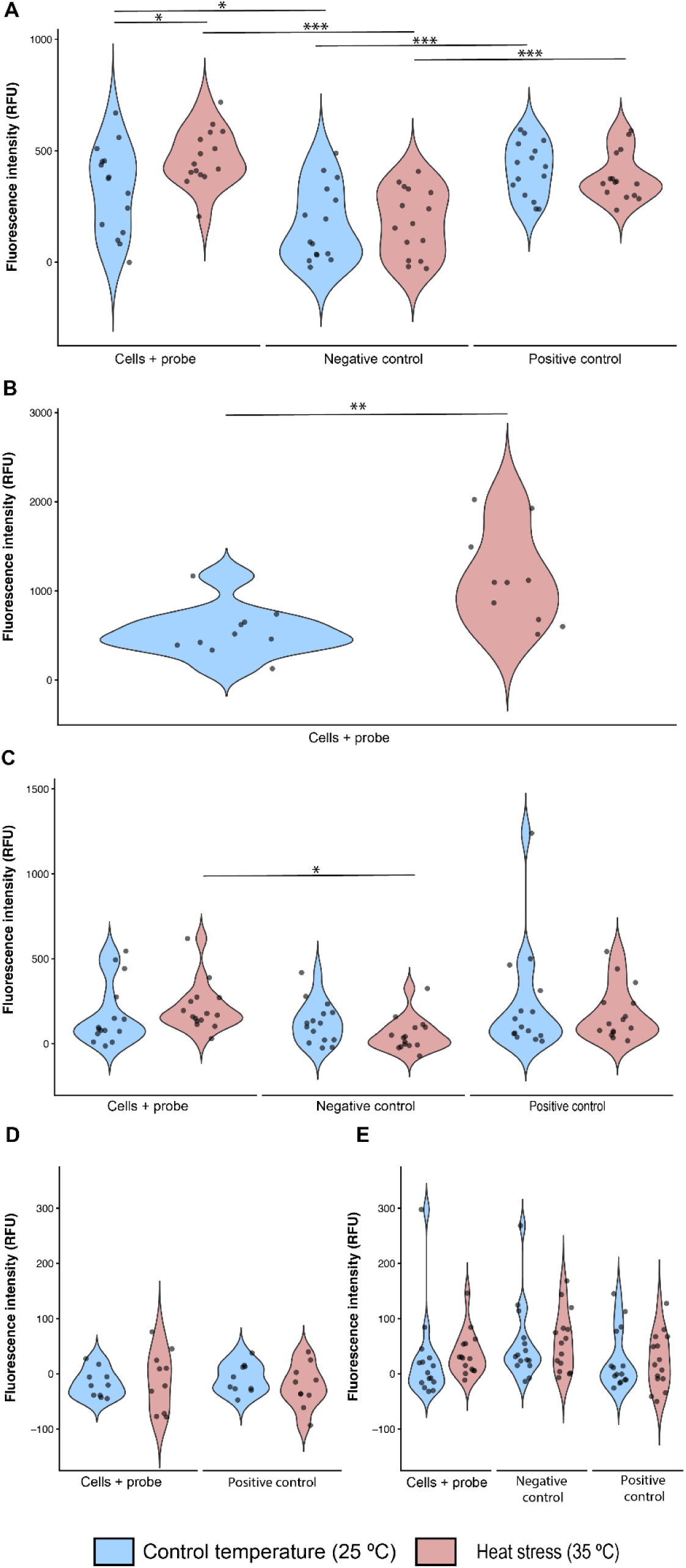
The average fluorescent intensity (RFU) of live coral cells stained with different fluorescent probes. **A)** Peroxynitrite-specific probe, (FRD, 10 µM) after 1 h incubation at 25 °C (control temperature, blue) or 35 °C (heat stress, red). Ebselen (EBS, 5.5 µM) was used as a negative control and 3-morpholinosydnonimine (SIN-1, 500 µM) was used as a positive control. Fluorescent intensity is corrected for background (unstained cells). Significance is indicated as * (p < 0.05) and *** (p < 0.001), ANOVA, Tukey’s test. **B)** General ROS probe, (carboxy DCFDA-AM, 20 µM) after 1 h incubation at 25 °C (control temperature, blue) or 33 °C (heat stress, red). Fluorescent intensity is corrected for background (unstained cells). Significance is indicated as ** (p < 0.01), Wilcox. **C)** Nitric oxide-specific probe, (DAF-2DA, 10 µM) after 1 h incubation at 25 °C (control temperature, blue) or 35 °C (heat stress, red). Luteolin (LUT, 33 µM) was used as a negative control and sodium nitroprusside (SNP, 0.1 mM) was used as a positive control. Fluorescence intensity is corrected for background (unstained cells). Significance is indicated as * (p < 0.05) and ** (p < 0.01) ANOVA, Tukey’s test. **D)** Superoxide (O ^-^)-specific probe, (MitoSOX Red, 0.25 µM) after 1 h incubation at 25 °C (control temperature, blue) or 35 °C (heat stress, red). 3-morpholinosydnonimine (SIN-1, 500 µM) was used as a positive control. Fluorescent intensity is corrected for background (unstained cells). No significance indicated, ANOVA, Tukey’s test. **E)** Superoxide dismutase (SOD, 16.67 U/µL) and MitoTempo (MTPO, 0.1 mM) were used as negative controls. Fluorescent intensity is corrected for background (unstained cells). No significance indicated, ANOVA, Tukey’s test.

**Figure 3.**
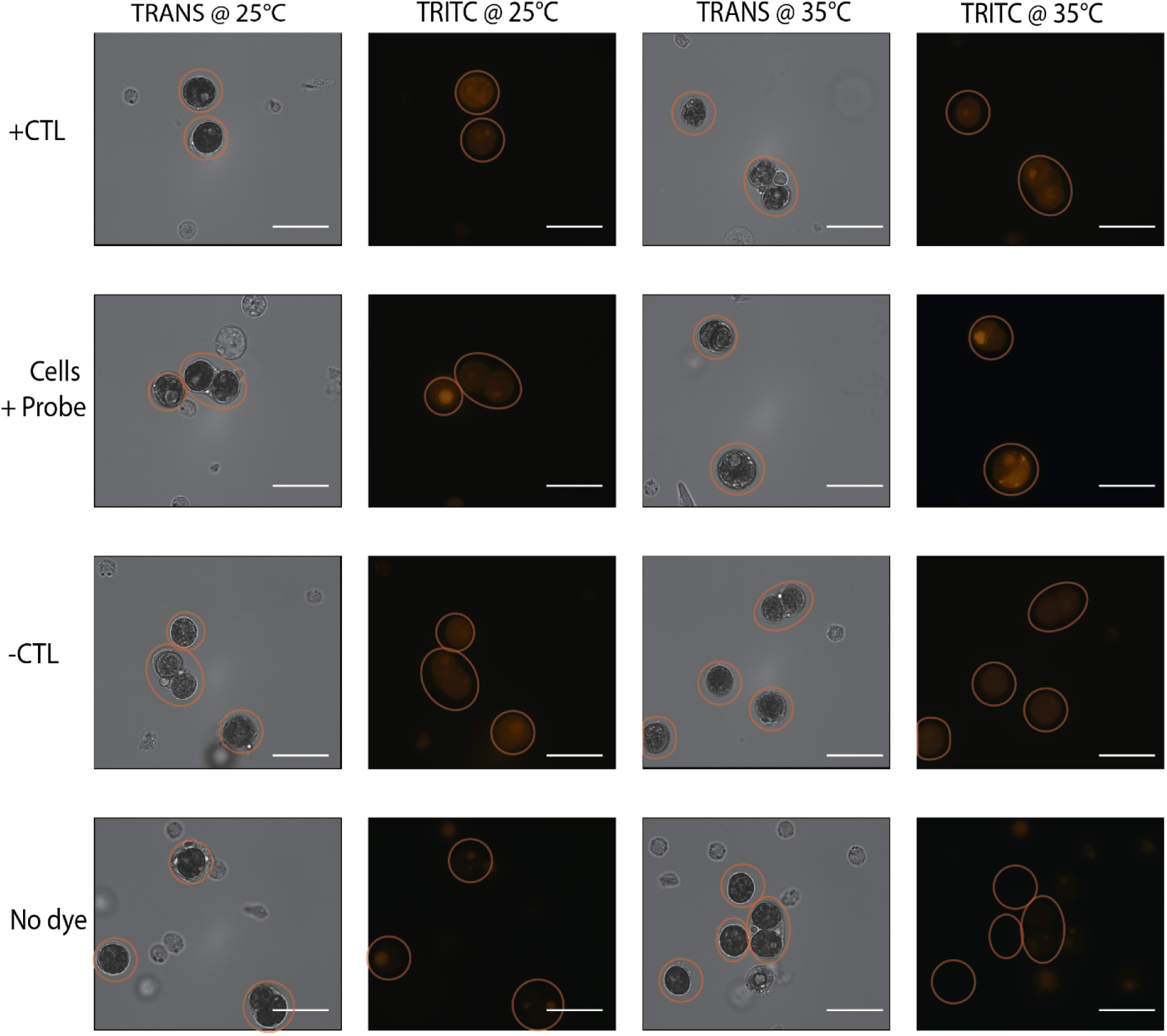
Visualized fluorescence (transmitted light (TRANS) and TRITC (λex/em 532–554/570–613 nm)) of live *P. acuta* cell suspension stained with FRD, a peroxynitrite probe (λex/em 600/638 nm, 10 µM), positive control SIN-1 (500 µM) (+CTL), and negative control ebselen (5.5 µM) (-CTL). Circles in micrographs depict the location of the symbiocyte. Scale bar = 20 *μ*m.

At 35 °C the average relative fluorescence intensity of cells treated with EBS (negative control) was 181.02 ± 10.06 RFU at 35 °C, a significant decrease (p < 0.001, Tukey’s test) compared to our unknown. This effect was also observed at control temperature (25 °C), where the negative control averaged 171.09 ± 11.30 RFU, though the difference was less significant (p < 0.05, Tukey’s test). The negative control was also significantly less fluorescent than the positive control at both temperatures (p < 0.001, Tukey’s test).

### ROS vs Peroxynitrite

As a result of heat stress, the average relative fluorescence intensity increased from 544.25 ± 28.01 RFU at 25 °C to 1141.48 ± 52.7 RFU at 35 °C, a significant increase in ROS production (p < 0.01, **Figure 2B**). The thermally stressed cells also showed a higher relative fluorescence intensity variability compared to control, where data were more similar.

### Peroxynitrite Precursors

#### Nitric oxide

The average relative fluorescence intensity of cells loaded with DAF2-DA was 168.47 ± 12.19 RFU at 25 °C and 211.43 ± 9.47 RFU at 35 °C. There was no significant increase in DAF-2DA fluorescence intensity as a result of thermal stress (**Figure 2C**), suggesting that nitric oxide levels were not detectably elevated in coral cells during thermal stress. The average relative fluorescence intensity of positive control was 231.14 ± 21.24 RFU at 25 °C and 177.31 ± 10.55 RFU at 35 °C, with no significant difference from the unknown samples (**Figure 2C**). However, a faint but observable fluorescent response was seen in positive control micrographs at both heat stress (35 °C) and control (25 °C) conditions, and in unknown samples at 25 °C (**Figure 4**).

**Figure 4.**
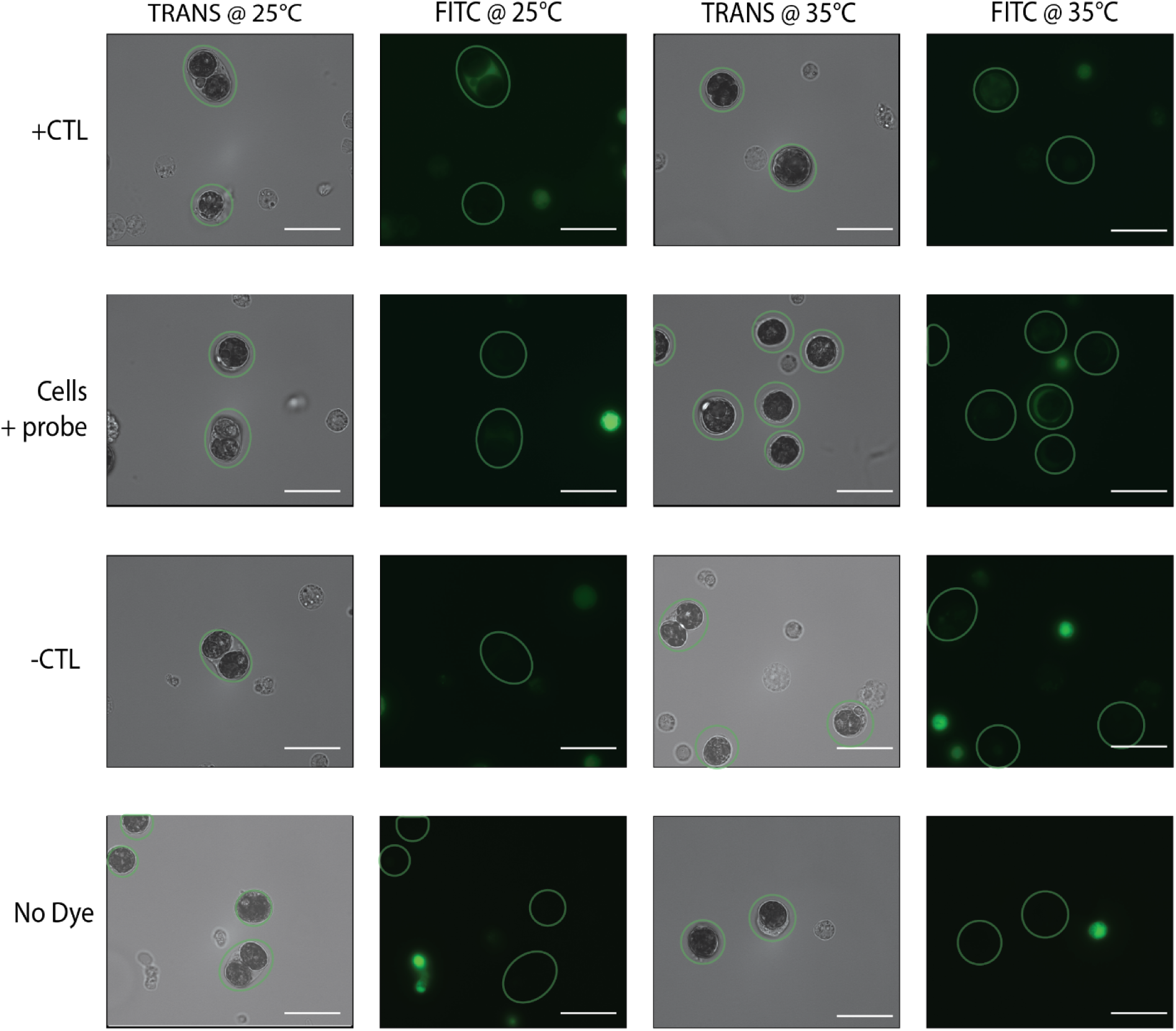
Visualize fluorescence (transmitted light (TRANS) and FITC (λex/em 475–490/515–540 nm))of live *P. acuta* cell suspension stained with DAF-2DA, a nitric oxide probe (λex/em 495/515 nm, 10 µM), positive control SNP (0.1 mM) (+CTL), and negative control LUT (33 µM) (-CTL 1). Circles in micrographs depict the location of the symbiocyte. Scale bar = 20 *μ*m.

DAF-2DA-specific fluorescence was typically seen within the coral host and symbiosome, while fluorescence within the algal symbiont was uncommon. Fluorescence within the coral host and symbiosome was not due to green fluorescent protein (GFP), as symbiocytes do not exhibit GFP fluorescence. At 35 °C the average relative fluorescence intensity of the negative control (luteolin) was 52.94 ± 6.48 RFU, a significant decrease (p < 0.01, Tukey’s test) relative to the cell-only treatment. Luteolin is reported as a nitric oxide scavenger (43); these data indicate that luteolin effectively scavenges the nitric oxide produced by the coral symbiocytes under thermal stress. The negative control was also significantly less fluorescent (p < 0.05, Tukey’s test) than the positive controls under thermal stress.

#### Superoxide

The average relative fluorescence intensity for unknown from positive and negative control trials was -17.19 ± 2.54 RFU and 25.41 ± 5.44 RFU, respectively at 25 °C. The corresponding average relative fluorescence intensity at 35 °C were -11.44 ± 5.38 RFU and 36.33 ± 2.68 RFU, respectively; neither showed a significant increase in fluorescence intensity as a result of thermal stress (**Figure 2D, E**). SIN-1, a source of superoxide, was trialled as a positive control but resulted in an average relative fluorescence intensity of -8.06 ± 2.68 RFU at 25 °C and -22.29 ± 3.96 RFU at 35 °C, showing no increase in fluorescence at either control or thermal stress conditions (**Figure 2D**). Two negative controls were trialled, SOD (25.16 ± 3.54 RFU at 25 °C and 23.09 ± 3.35 RFU at 35 °C) and MTPO (54.58 ± 4.71 RFU at 25 °C and 23.09 ± 3.35 RFU at 35 °C), but neither could elicit any significant change in fluorescence intensity (**Figure 2E**). No visible response was observed in microscope images (**Figure S2**).

### Time-series assay (DAF-2DA, MitoSOX, carboxy-DCFDA-AM)

*P. acuta* cell suspensions treated with MitoSOX Red exhibited a modest increase in fluorescence over time, which can be accelerated by adding hydrogen peroxide (3.3 µM, **Figure 5A**). However, introducing thermal stress did not increase fluorescence production.

**Figure 5.**
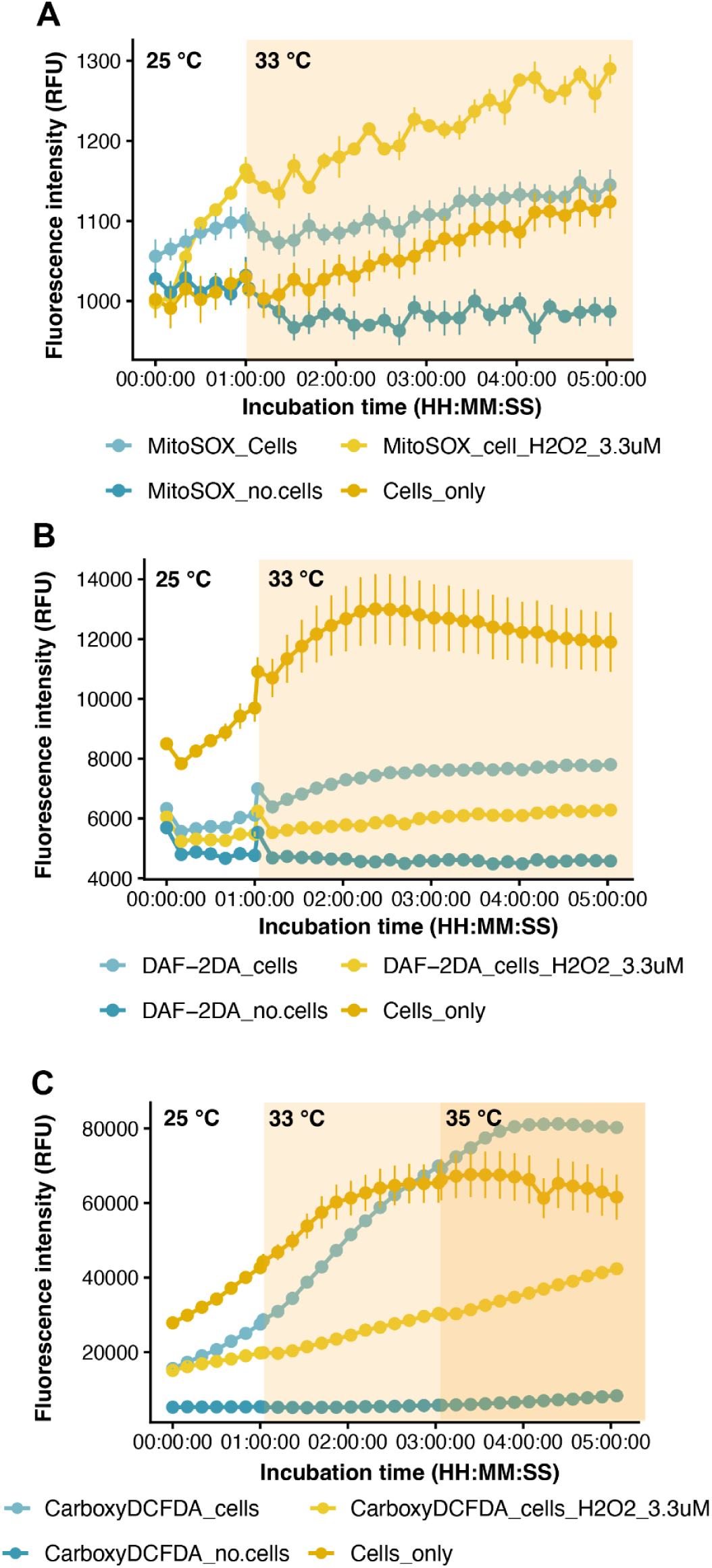
Time series showing fluorescence of *P. acuta* cell suspensions during exposure to thermal stress: **A)** cells stained with superoxide probe (MitoSOX_Cells), including a hydrogen peroxide-spiked control (MitoSOX_cell_H2O2_3.3uM), a dye-only control (MitoSOX_no cells) and an unstained cell-only control (Cells_only); **B)** cells stained with nitric oxide probe (DAF-2DA), including a hydrogen peroxide-spiked control (DAF-2DA_cell_H2O2_3.3uM), a dye-only control (DAF-2DA_no cells) and an unstained cell-only control (Cells_only); **C)** cells stained with superoxide probe (CarboxyDCFDA_cells), including a hydrogen peroxide-spiked control (CarboxyDCFDA_cell_H2O2_3.3uM), a dye-only control (CarboxyDCFDA_no cells) and an unstained cell-only control (Cells_only). Error bars are equal to the standard error of each data point.

The addition of DAF-2DA and carboxy-DCFDA-AM to *P. acuta* cell suspensions decreased their fluorescence (**Figure 5B**, **C**) that remains unexplainable. Moving baselines present a further challenge: unstained coral cell suspensions show an increase in fluorescence when thermal stress is applied (**Figure 5B**, **C**). The addition of hydrogen peroxide (3.3 µM) was also associated with a decrease in fluorescence (**Figure 5A**, **C**), contrary to expectation, particularly for carboxy-DCFDA-AM which is a broad-spectrum ROS dye. Lower concentrations of hydrogen peroxide (1.7, 0.8 0.4 µM, data not shown) produced an identical decrease in fluorescence for both DAF-2DA- and carboxy-DCFDA-AM-loaded coral cells.

Following this initial decrease, both DAF-2DA (**Figure 5B**) and carboxy-DCFDA-AM (**Figure 5C**) exhibited an increase in fluorescence intensity over time, followed by a plateau at 2 h and 4 h, respectively. Under thermal stress, the rate of production of nitric oxide (DAF-2DA) and ROS (carboxy-DCFH-DA) production increased in the first hour by factors of 9 and 2, respectively.

## Discussion

### Peroxynitrite pathway

The significant difference in FRD relative fluorescence intensity observed in *P. acuta* cells exposed to thermal stress (**Figure 2**) implies an uptick in RNS production as a result of heat stress. Compared with nitric oxide (**Figure 5**) and superoxide (**Figure 7**) trials, peroxynitrite returned significant differences across a greater number of treatment comparisons. This supports the hypothesis that the nitric oxide and superoxide probes did not show significant changes due to thermal stress because the targeted radicals react with each other at an incredibly fast rate (2 × 10^10^ M^-1^ s^-1^ (48)), up to ten times faster than the reaction rate of superoxide with superoxide dismutase (2 × 10^9^ M^-1^ s^-1^ (49)). This suggests that as nitric oxide is being produced as a response to thermal stress, so is superoxide and this co-production prevents the accumulation of the two separate radicals; instead, the two radicals react with each other to form peroxynitrite. By contrast, DAF-2DA fluorescence did increase in response to thermal stress in time-series experiments (**Figure 5**), matching the trend seen for broad-spectrum ROS; however, this increase may instead reflect a thermal stress response specific to the green fluorescent protein phenotype of the genet used in this study (50), in which pigment proteins are upregulated under thermal stress as a photoprotective mechanism, rather than a direct measure of nitric oxide production. This remains a plausible interpretation and would need to be confirmed using a differently-pigmented genet as a comparison.

Mechanistically, the presence of peroxynitrite implies that nitric oxide and superoxide must be present within the system. The performance of the DAF-2DA nitric oxide probe and the MitoSOX superoxide probe may be hampered by the seawater-based media, or by the multiple membrane barriers within the symbiocyte (**Section 4.2**). It is also possible that the RNS produced are quenched by the cells’ endogenous antioxidant system, or transformed into downstream reactive species such as peroxynitrite (5,16), before it is able to react with the probe.

Detecting each reactive species individually allows us to piece together the pathways of nitrosative stress. This non-model system does not lend itself to direct application of established mammalian tools and techniques, therefore it is important to build redundancy into the experimental design. Measuring only superoxide or nitric oxide may lead to an incorrect conclusion; that the nitrosative pathway is insignificant in the context of thermally-induced oxidative stress. By detecting an increase in peroxynitrite concentration in live, thermally stressed *P. acuta* cells, our results are consistent with activation of the nitrosative pathway under thermal stress. Due to the cytotoxicity of peroxynitrite (15), the nitrosative stress pathway may play a role in the breakdown of symbiosis, independently of oxidative stress. Similarly, recent single-cell based research found that thermal stress did not have a significant enough effect on ROS production to trigger the bleaching cascade at the cellular level (26). However, they did report an increase in endosymbiont ROS production under thermal stress using analogous broad-spectrum dyes, a result consistent with our own thermal stress trials using carboxy-DCFDA-AM (**Figure 2**). In our data, ROS production increased by a greater factor than peroxynitrite production as a result of thermal stress, highlighting that nitrosative stress could be obscured without a species-specific dye.

### Method development and limitations

Selecting a buffer system that preserved *P. acuta* cell health without impeding probe performance was a central challenge. Most fluorescent probes used here, including the peroxynitrite-specific PNP-1 (47), are optimised for mammalian systems (e.g., 1X PBS), which differ from seawater in both pH (7.4 vs. 8.0–8.3) and salinity (9‰ vs. 35‰); deviation from these mammalian parameters compromised probe sensitivity and selectivity (**Figure S1**), while deviation from marine conditions reduced cell viability. Standard coral cell culture media (ASW, DMEM, FBS, gentamicin, and anti-anti; (25)) was similarly unsuitable for reactive species detection (**Figure S1**), and unsupplemented seawater introduced dissolved metals that scavenge radical species. Seawater supplemented with EDTA (a metal chelator, 20 µM) offered a reasonable compromise, consistent with established concentrations (51).

Positive and negative controls posed further challenges. SIN-1 was an ineffective upregulator of peroxynitrite (**Figure 2**); although SIN-1 is thought to function in seawater (52) and has previously served as a positive control for FRD (29), peroxynitrite is markedly less stable in seawater (70 ms lifetime, (53) than in mammalian culture media (800–1000 ms, (15)), likely limiting the amount reaching intracellular FRD before decomposing. As a result, our evidence for peroxynitrite detection rests on a thermal-stress-associated increase in FRD fluorescence together with an effective negative control (ebselen); we did not obtain an independent, in-system positive control confirming that FRD responds to exogenous peroxynitrite under our specific buffer and live-cell conditions. We consider this an acceptable, if incomplete, control structure given the mechanistic explanation for SIN-1’s failure described above, though direct validation of FRD responsiveness to spiked peroxynitrite in live *P. acuta* cell suspensions would strengthen this result and represents an important target for future work. By contrast, ebselen, a peroxynitrite scavenger (29), was an effective negative control under thermal stress. SNP failed as a positive control for nitric oxide (**Figure 2**), consistent with existing literature on Symbiodineceae (54) despite prior success in symbiotic anemones (21), while the negative control (luteolin) reduced fluorescence under thermal stress as expected. Neither the positive (SIN-1) nor negative (MTPO, SOD) controls for superoxide performed effectively (**Figure 2**). In the case of SIN-1, it may be that the reaction of superoxide and nitric oxide outpaces the reaction of superoxide with the MitoSOX probe (55).

Probe performance was likely constrained by the multiple membrane barriers within the symbiocyte (27) and by the salinity and alkalinity of ASW-based buffers (27,34); FRD was a notable exception, remaining stable across pH 6–9 (29), which supports its use as a robust platform for peroxynitrite detection. Where dyes underperformed, three explanations are plausible: the target free radical was genuinely absent or below detectable levels (defensible only where the positive control confirmed the dye’s ability to detect it); a reaction or decomposition rate too fast for the dye to capture; or non-model buffer conditions preventing the dye from reacting altogether. DAF-2DA fluorescence, for instance, was detected only in the coral host and symbiosome, not the algal symbiont (**Figure 4**), suggesting the polar, deacetylated DAF-2 product forms in the host but cannot cross further membranes to reach the symbiont. MitoSOX Red, while free of autofluorescence interference, may also be unsuitable for superoxide detection above pH 7.4 (56); alternative superoxide-detection methods (57) should be explored in future coral work.

Time-series measurements were further complicated by endogenous GFP fluorescence (λex/em = 478–508/500–520 nm; (30)) specific to the green phenotype of *P. acuta* used in this study, which overlapped with DAF-2DA and carboxy-DCFDA-AM emission and could not be reliably disentangled from probe signal (**Figure 5B, C**); both probes showed an unexplained decrease in fluorescence upon addition to cell suspensions, further reduced by hydrogen peroxide addition. The mechanism behind this unexpected decrease is not resolved by the present data, but two possibilities warrant consideration. First, DAF-2 and DCF are both fluorescein-based fluorophores, which are susceptible to oxidative photobleaching at sufficiently high oxidant exposure; give the multi-hour incubation used in the time-series assays, cumulative oxidative degradation of the fluorophore itself may outpace probe activation. Second, the cell culture medium’s fetal bovine serum component carries endogenous catalase and peroxidase activity, which may substantially degrade extracellular hydrogen peroxide before it reaches the intracellular esterase-activated probe (38). Distinguishing between these, and other, possible explanations will require dedicated follow-up experiments. Taken together, these caveats mean that firm conclusions about nitric oxide, superoxide, and general ROS production cannot be drawn from the time-series data alone, underscoring the importance of appropriate controls and careful interpretation when applying these tools to non-model systems.

The use of a single monoclonal genet allowed us to isolate probe and assay performance from the substantial genotype-driven variation in thermal tolerance and bleaching susceptibility known in *P. acuta* (39), which would otherwise have confounded interpretation of a first attempt at live-cell peroxynitrite detection. However, this design means our results reflect the redox response of a single genetic background, and the extent to which the reported thermal-stress-associated peroxynitrite increase generalizes across genotypes, or across coral species more broadly, remains untested. Future work applying this assay platform across multiple genets and species will be needed to establish the generality of this response.

## Conclusions

In conclusion, these results show that thermal stress is associated with an increase in intracellular peroxynitrite concentration in live *P. acuta* cells. Its precursors, nitric oxide and superoxide, showed no corresponding increase, a result likely reflecting their rapid consumption in the reaction that forms peroxynitrite, rather than the absence of production, and highlight the difficulty of resolving fast-reacting radicals with fluorescent probes in non-model systems. Despite the methodological challenges above, the detection of peroxynitrite in live, thermally stressed *P. acuta* cells represents a step forward in understanding the mechanism of coral bleaching, and to our knowledge, the first such detection in a live coral cell system. Combining fluorescent probes with our cell culture media platform provides a viable, if still imperfect, platform for studying oxidative and nitrosative stress pathways in live coral cells under thermal stress. Future work should continue to optimize probe performance within non-model systems, and should prioritize linking intracellular peroxynitrite production to markers of coral cell health, which would help establish peroxynitrite’s role as a mediator of bleaching in coral symbiocytes.

## Acknowledgments

We thank Dr Heitzman for his preliminary work on several fluorescent probes. Although that data is not included in this manuscript, it helped inform the early stages of this research. This study was supported by the National Science Foundation (NSF) award numbers 2316389 and 2238563 and Arizona State University’s Presidential Strategic Initiative Fund (PI Roger). Research reported in this publication was supported by a Beckman Young Investigator Award from the Arnold and Mabel Beckman Foundation (D.W.D.) G.D.K was supported by this award.

