## Supplementary Information for "First detection of peroxynitrite in live coral cells during thermal stress"

**Table S1.** Materials used

| **Name (abbreviation)** | **Stock purity/concentration** | **λex/em** | **Cat. no** | **Supplier** | **Function/target** |
|---|---|---|---|---|---|
| Dulbecco's Modified Eagle Medium, without phenol red or sodium pyruvate | 1X | - | 21063029 | Gibco, NY, USA | Cell culture media (CCM) component |
| Fetal Bovine Serum | - | - | A5670701 | Thermo Fisher Scientific, MA, USA | CCM component |
| Antibiotic-antimycotic | 100X | - | 15240096 | Gibco, UK | CCM component |
| Gentamicin | 50 mg/mL | - | 15750060 | Gibco, UK | CCM component |
| EDTA | 0.5 M | - | AM9261 | Invitrogen, Vilnius, LTU | Metal scavenger |
| Coral Dip disinfectant | - | - | - | Coral Rx, FL, USA | Coral dip disinfectant |
| 4,5-Diaminofluorescein diacetate (DAF-2DA) | ≥ 98% | 495/515 nm | 251505-M | Sigma-Aldrich, MA, USA | Nitric oxide probe |

| | | | | | |
|---|---|---|---|---|---|
| Luteolin (LUT) | 3 mM, ≥90% | - | 440025 | Sigma-Aldrich, MA, USA | Nitric oxide inhibitor and scavenger |
| Sodium nitroprusside (SNP) | ≥99% | - | 71778 | Sigma-Aldrich, MA, USA | Induces nitric oxide production |
| BioTracker Far-Red Peroxynitrite Live Cell Dye (FRD) | ≥ 98% | 600/638 nm | SCT052 | Sigma-Aldrich, MA, USA | Peroxynitrite probe |
| Ebselen (EBS) | ≥ 98% | - | 5245 | Tocris, UK | Peroxynitrite scavenger |
| 3-Morpholinosydnonimine hydrochloride (SIN-1) | ≥ 97% | - | M5793-25MG | Sigma-Aldrich, MA, USA | Releases nitric oxide and superoxide to form peroxynitrite |
| MitoSOX Mitochondrial Superoxide Indicator Red (MitoSOX) | ≥ 80% | 396/610 nm | M36008 | Thermo Fisher Scientific, MA, USA | Superoxide probe |
| Superoxide dismutase (SOD) | 3000 U/mL | - | S5395 | Sigma-Aldrich, MA, USA | Superoxide antioxidant |
| MitoTEMPO (MTPO) | ≥ 98% | - | SML0737 | Sigma-Aldrich, MA, USA | Superoxide dismutase mimetic |

| | | | | | |
|---|---|---|---|---|---|
| 6-Carboxy-2’,7’-dichlorodihydrofluorescein diacetate, di(acetoxymethyl ester) (Carboxy DCFDA-AM) | ≥ 90% | ~492–495/517–527 nm | C2938 | Thermo Fisher Scientific, MA, USA | Broad spectrum ROS probe |
| Hydrogen peroxide | 35% w/w aqueous solution, stabilized | - | L14000.AP | Thermo Fisher Scientific, MA, USA | - |
| PNP-1 | - | 450/525 nm | - | Domaille group, Colorado School of Mines | Peroxynitrite probe |
| Peroxynitrite | 73.3 mM | - | 81565, Batch no. 0660628 | Cayman chemical, MI, USA | - |
| PBS, pH 7.4, without calcium chloride or magnesium chloride | 10X | - | 70011-044 | Gibco, UK | Buffer |

### Supplementary information

***S1.*** *Coral husbandry:*

A monoclonal population of *Pocillopora acuta* (green phenotype) specimens originating from Hawaii, propagated by and obtained from the Putnam Lab (University of Rhode Island, RI, USA) was maintained in a 170 L aquaria system with 24 L sump. Artificial seawater (ASW, 35 ‰ salinity; Tropic Marin Pro-Reef Salt) was maintained at 25 °C (600 W titanium aquarium heater with Inkbird ITC-306 A InkBird temperature controller) with a 12 h:12 h photoperiod at 150–200 µmol photons $m^{-2}$ $s^{-1}$ irradiance (AP9x LED lights; Kessil). The ASW was constantly filtered (200 µm sock filter; Aquatic Experts), skimmed for dissolved organic compounds (Octo Classic 110S protein skimmer; Reef Octopus) and UV sterilized (Aqua Ultraviolet). Aquaria were cleaned once a week to further limit biofouling, and a 10% water change also performed weekly. The ASW parameters were monitored daily (pH, salinity, redox potential, temperature, carbonate hardness [kH]) or weekly (calcium, magnesium, phosphate, nitrate) using probes (Proflux 4 GHL) and test kits (API saltwater test kits, Salifert magnesium test kit). Buffers were dosed as necessary to adjust ASW parameters (Fauna Marin Balling Light Set; Fauna Marin). *P. acuta* were fed weekly with live *Artemia salina* nauplii (E-Z EGG; Brine Shrimp Direct).

***S2.*** *Cell viability*

Hoechst 33342 (cell permeable DNA stain, λex/em = 350/461 nm, f.c. 20 µM, Thermo Scientific, Cat. no. 62249) and SYTOX orange (dead cell stain, λex/em = 547/570 nm, f.c. 2.5 µM, Invitrogen, Cat. no. S34861) were added to an aliquot of cell suspension in CCM before incubation (30 min, 25 °C, dark (illuminated incubator, samples wrapped in aluminum foil)). Post incubation, the suspension was washed and resuspended in CCM and added to a hemocytometer. Viability analysis was performed in a microplate reader (Agilent BioTek Cytation 5) using the Cellular Analysis function in BioTek Gen 5 software. Cell counts were performed using DAPI (λex/em = 377/447 nm) (total nucleus count) and TRITC (λex/em = 556/600 nm) images, with detection settings: intensity maximum = 11000, object size 2–13 µm, plug size 1000 µm × 1000 µm. Cell viability (live cells as a proportion of total cell count) was calculated for five plugs using Equation 1, and the average taken to return a representative cell viability. For assays, cell suspensions were centrifuged (805 × *g*, 3 min, 25 °C) and the cell pellet resuspended in ASWwith EDTA at 1 × $10^6$ live cells/mL.

**Equation 1:** $Cell\ viability\ (\%) = \frac{live\ cell\ count\ (DAPI\ count - TRITC\ count)}{DAPI\ count} \times 100$

***S3.*** *Nitric oxide imaging:*

To detect nitric oxide production, *P. acuta* samples (1 × $10^6$ live cells/mL) were stained with DAF-2DA (f.c. 10 µM) and incubated (30 min, 25 °C, dark) before washing and resuspending in ASW with EDTA. Samples were then subject to their assigned thermal conditions (1 h at 25 °C or 35 °C). Sodium nitroprusside (SNP, f.c. 0.1 mM) and luteolin (LUT, f.c. 33 µM) were dosed into their respective treatment tubes. All samples were incubated at their respective thermal conditions for a further 30 min before washing and resuspending the positive and negative controls in ASW with EDTA before imaging.

***S4.*** *Superoxide imaging:*

To detect mitochondrial superoxide production, MitoSOX working solutions (0.5 µM) were prepared from MitoSOX stock solution (5 mM) and added to *P. acuta* samples (1 × $10^6$ live cells/mL) at a 1:1 ratio (f.c. 0.25 µM). The samples were incubated (30 min, 25 °C, dark) before washing and resuspending in ASW with EDTA. 3-Morpholinosydnonimine hydrochloride (SIN-1, f.c. 500 µM), superoxide dismutase (SOD, f.c. 16.67 U/µL) and MitoTempo (MTPO, f.c. 0.1 mM) were dosed to their respective cell suspensions before further incubation under respective thermal conditions (30 min, 25 or 35 °C). Positive and negative controls were centrifuged and resuspended in ASW with EDTA before resting in their respective thermal conditions (25 °C or 35 °C) until imaging.

***S5.** Peroxynitrite imaging:*

To detect peroxynitrite production, *P. acuta* negative control samples (1 × $10^6$ live cells/mL) were pretreated with ebselen (f.c. 5.5 µM) and incubated (30 min, 25 °C, with light) while dye only and positive control samples were stained with far red dye (FRD, f.c. 10 µM) and incubated (30 min, 25 °C, dark). All samples (dye only, positive, and negative) were rinsed and resuspended in ASW with EDTA. Negative control samples were then stained with FRD (f.c. 10 µM) and incubated (30 min, 25 °C, dark). After incubating with FRD, positive and negative controls were dosed with SIN-1 (f.c. 500 µM). All samples were then subjected to continued thermal stress (1 h at 25 °C or 35 °C, light). After the final incubation, the positive and negative controls were washed and resuspended in ASW with EDTA before imaging.

***S6.** Broad spectrum ROS imaging:*

*P. acuta* aliquots (1 × $10^6$ live cells/mL) were treated with carboxy-DCFDA-AM (f.c. 20 µM) and incubated (30 min, 25 °C, dark) before washing and resuspending the sample in ASW with EDTA. Samples were subject to respective thermal stress (1 h at 25 °C or 33 °C, light), washed and resuspended in ASW+EDTA before imaging.

***S7.** Nitric oxide time-series:*

DAF-2DA (f.c. 10 µM) was added to *P. acuta* cell suspension (1.0 × $10^6$ live cells $mL^{-1}$ in CCM). The samples were incubated (15 min, 25 °C, dark), centrifuged and resuspended in CCM and transferred to a black 96-well plate (100 µL/well) for analysis. Hydrogen peroxide (f.c. 3.3

μM) was added to select wells as a positive control. A sample consisting of CCM only was stained with DAF-2DA and used as a method blank. Samples were incubated for 1 h at 25 °C and 4 h at 33 °C in the dark, and fluorescence was measured at λex/em = 495/515 nm every 10 min using a microplate reader and imager.

***S8.*** *Superoxide time-series:*

DMSO was added to one vial of MitoSOX red (f.c. 5 mM), diluted to 0.5 μM in CCM. The working MitoSOX solution (f.c. 0.25 μM) was added to *P. acuta* cell suspension ($1.0 \times 10^6$ live cells $mL^{-1}$ in CCM) in a 1:1 ratio and incubated (30 min, 25 °C). Cells with MitoSOX were then washed and resuspended in CCM, and plated in a black 96-well plate for analysis. Hydrogen peroxide was added to select wells as a positive control (f.c. 3.3 μM). CCM was stained with MitoSOX and used as a method blank. Samples were incubated for 1 h at 25 °C followed by 4 h at 33 °C, and fluorescence was measured at λex/em = 396/610 nm every 10 min using a microplate reader and imager.

***S9.*** *Broad spectrum ROS time-series:*

Carboxy-DCFDA-AM was added to *P. acuta* cell suspension ( $1.0 \times 10^6$ live cells $mL^{-1}$ in CCM) to a concentration of 20 μM and incubated for 30 min at 25 °C. The cells were then washed and resuspended in CCM, and plated in a black 96-well plate to rest for 15 min before analysis. Hydrogen peroxide was added to select wells as a positive control (f.c. 3.3 μM). A sample consisting of CCM only was stained with carboxy-DCFDA-AM and used as a method blank. Samples were incubated in the plate reader for 1 h at 25 °C, then 2 h at 33 °C, then 2 h at 35 °C, and fluorescence measured at λex/em = 495/515 nm every 10 min in a microplate reader and imager.

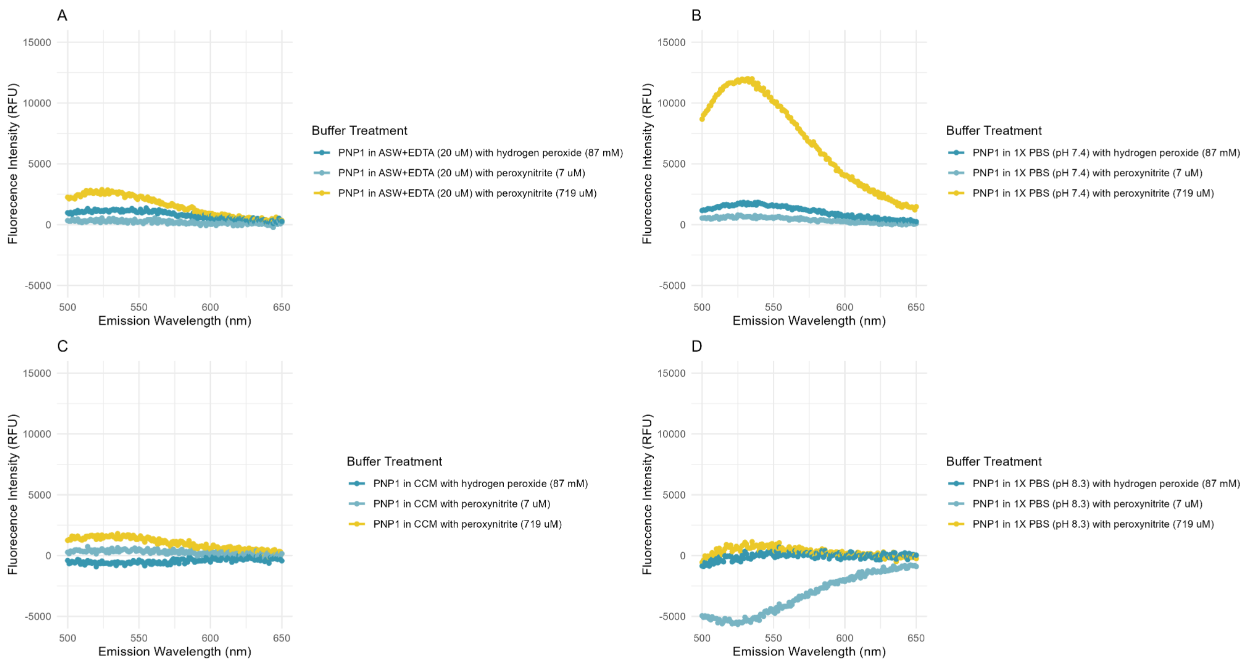


**Figure S1.** Fluorescence spectra (ex = 450 nm) for PNP-1, a peroxynitrite probe, in **A)** ASW+EDTA (20 µM), **B)** 1X PBS (pH 7.4), **C)** cell culture media (CCM), and **D)** 1X PBS (pH 8.3). Spectra are corrected for baseline fluorescence (buffer+PNP-1). Buffers are spiked with peroxynitrite (719, 7 µM) and hydrogen peroxide (87 mM) to assess PNP-1 sensitivity and selectivity.

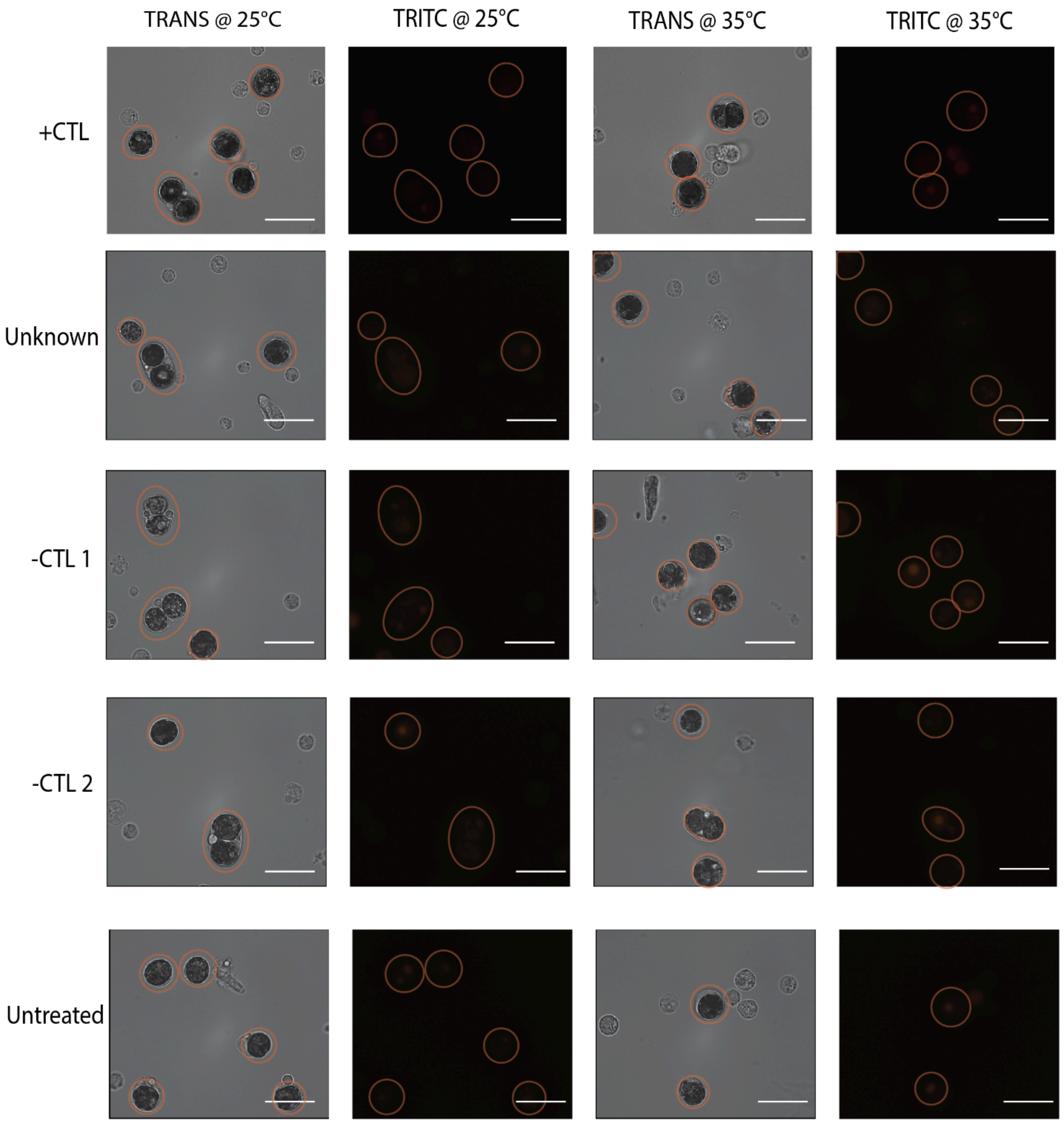


**Figure S2** Visualized fluorescence (transmitted light (TRANS) and TRITC (λex/em 532–554/570–613 nm)) of live *P. acuta* cell suspension stained with MitoSOX Red, a superoxide probe (λex/em 400/590 nm, 0.25 μM), positive control SIN-1 (500 μM) (+CTL), negative control MitoTempo (0.1 mM) (-CTL 1), and negative control superoxide dismutase (16.67 U/μL) (-CTL 2). Circles in micrographs depict the location of the symbiocyte. Scale bar = 20 $\mu$m.
